# Evaluation of the Accelerate Pheno™ system to identify bacteria and determine antimicrobial susceptibility in positive blood cultures

**DOI:** 10.1101/621268

**Authors:** Mariana J. Fernandez-Pittol, Javier Morales, Elisa Rubio, Assumpta Fasanella, Izaskun Alejo-Cancho, Manel Almela, Francesc Marco, Jordi Vila, Climent Casals-Pascual

## Abstract

**Introduction:** New platforms have recently been developed to reduce response time of identification and antimicrobial susceptibility of bacterial isolates in positive blood cultures from patients with bloodstream infections. The Accelerate Pheno™ system (Accelerate Diagnostics, Inc.) provides information on pathogen identification and antibiotic susceptibility in approximately 1.5 and 7 hours, respectively.

**Methods:** In this study we compared the Accelerate Pheno™ system with the standard procedure used in our laboratory. A total of 41 blood cultures were prospectively analysed with the Accelerate Pheno™ system and our standard methods, which include identification by MALDI-TOF mass spectrometry and antibiotic susceptibility testing (AST) by BD Phoenix system and E-test.

**Results:** The correlation between the two methods using Cohen’s kappa coefficient was 0.82; mean (sd) time of identification for MALDI-TOF MS was 0.7 (0.22) hours and 1.43 (0.14) hours for the Accelerate Pheno™ system. The mean (sd) time of AST with the BD Phoenix system was 15.85 (2.57) hours and with the Accelerate Pheno™ system 6.7 (0.12) hours. AST results showed an overall essential agreement of 92% for the minimal inhibitory concentrations (MIC) and an overall category agreement of 96%. Among Gram positive isolates, essential and category agreements of 100% were observed. In Gram negative isolates 10 discrepancies were detected, which were classified as 7 major and 3 minor errors. Discrepancies in the Accelerate Pheno™ system were observed particularly for *P. aeruginosa.*

**Conclusion:** The Accelerate Pheno™ system can improve turn-around time in the management of patients with bloodstream infections.

## Introduction

The management of patients with positive blood culture requires prompt identification and antimicrobial susceptibility testing (AST) since the time to response is critical to implement an adequate antibiotic treatment and to reduce mortality. New platforms are being developed to provide targeted therapy and thus decrease morbidity and mortality rates in sepsis (1–4). The Accelerate Pheno™ system (Accelerate Diagnostics, Inc.) is an automated platform that uses single-cell microbiology analysis technology to rapidly identify pathogens (1.5 hours) and to determine antibiotic susceptibility (7 hours) directly from the positive blood culture bottle (5–7). The system combines gel electrofiltration and fluorescence in situ hybridization for bacterial identification, as well as automated microscopy to analyse bacterial growth rates and for extrapolating MIC values (5, 6). This study prospectively compares the Accelerate Pheno™ system with the matrix-assisted laser desorption ionization–time of flight mass spectrometry (MALDI-TOF MS, Bruker) for identification and the BD Phoenix system (Becton Dickinson, New Jersey, U.S) for AST.

## Methods

The study was designed as a method validation. All laboratory samples were anonymized and unlinked from the hospital database. Only laboratory parameters were recorded and used for this study. No personal or clinical information from patients was used in the study. A total of 41 positive blood cultures were analysed. According to the initial Gram staining, this series contained 18 Gram-negative rods, 21 Gram-positive cocci (9 *Staphylococcus* spp and 12 *Streptococcus* spp) and 2 yeasts. The standard procedure used in our laboratory includes direct identification based on the MALDI-TOF MS (Bruker), and antimicrobial susceptibility testing (AST) performed with the BD Phoenix system (Becton Dickinson, New Jersey, U.S) for Gram-negative rods, *Staphyloccus aureus* and *Enterococcus* spp. This panel includes information of 34 antibiotics in Gram-positive (*Staphyloccus aureus*) and 31 en Gram-negative. The Accelerate Pheno™ system panel provides information of 8 antibiotic for Gram-positive and 15 antibiotics for Gram-negative. When S. aureus infection was suspected or the MALDI-TOF MS identified *S. aureus*, an automated PCR system GenomEra (GenomEra, Alere) was also performed for confirmation (*S. aureus* and coagulase negative staphylococci) and to detect methicillin resistance (detection of the mecA gene). The identification (ID) and AST of the Accelerate Pheno™ system were performed side by side with our standard microbiological procedure. AST discrepancies between the two methods were resolved using the E-test method.

## Results

Table 1 shows the organisms identified using both systems. Although the correlation between the two methods was not perfect, there was a good agreement in terms of microorganism identification (Cohen’s kappa coefficient =0.82). The Accelerate Pheno™ system produced three false identifications. In the first case, Accelerate Pheno™ falsely identified *S. pneumoniae* plus *Streptococcus* spp in a sample where MALDI-TOF MS showed no identification and the subculture yielded Coagulase negative staphylococci (CNE) and viridians streptococci. In a second case, *S. agalactiae* plus *Streptococcus* spp were detected, where the MALDITOF and subculture only identified *S. agalactiae*. Finally, *E. faecalis* plus *P.aeruginosa* were identified by Accelerate Pheno™ system, where MALDI-TOF MS and subculture only detected *E. faecalis*. In two cases the results of the Accelerate Pheno™ system did not correlate with results from the MALDI-TOF MS. In one instance the Accelerate Pheno™ system identified three microorganisms (*P.aeruginos*a, *C.glabrata* and *S.aureus*) whereas MALDITOF MS detected only *P.aeruginosa* which was further confirmed by culture. In a second instance, MALDI identified *P. aeruginosa* but no identification was provided by Accelerate Pheno™. Direct MALDI-TOF MS was not performed in two yeast positive blood cultures due to low inoculum. In both cases the Accelerate PhenoTM system could not identify these microorganisms (by culture: *C. albicans* and *C. tropicalis*). The mean (SD) times of ID for MALDI-TOF MS and for the Accelerate PhenoTM system were 0.7 (0.22) hours and 1.43 (0.14) hours, respectively. The mean (SD) times to AST result with the BD Phoenix system (Becton Dickinson) and with the Accelerate Pheno™ system were 15.85 (2.57) hours and 6.7 (0.12) hours, respectively (Figure 1).

**Table 1.**
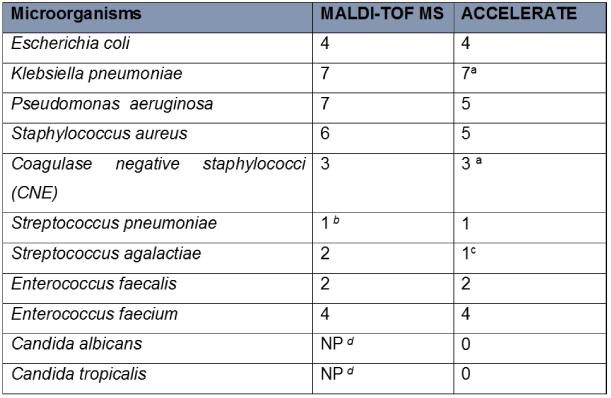
Comparison of microorganisms Identified using Accelerate and MALDITOF. ^a^ Accelerate only identified the genus level (*Klebsiella* spp and coagulase negative staphylococci). ^b^ Confirmed with the agglutination test for direct detection of capsular antigens. ^c^ *S.agalactiae* plus *Streptococus* spp. ^d^ Not performed. Identification obtained from culture plates.

**Fig. 1.**
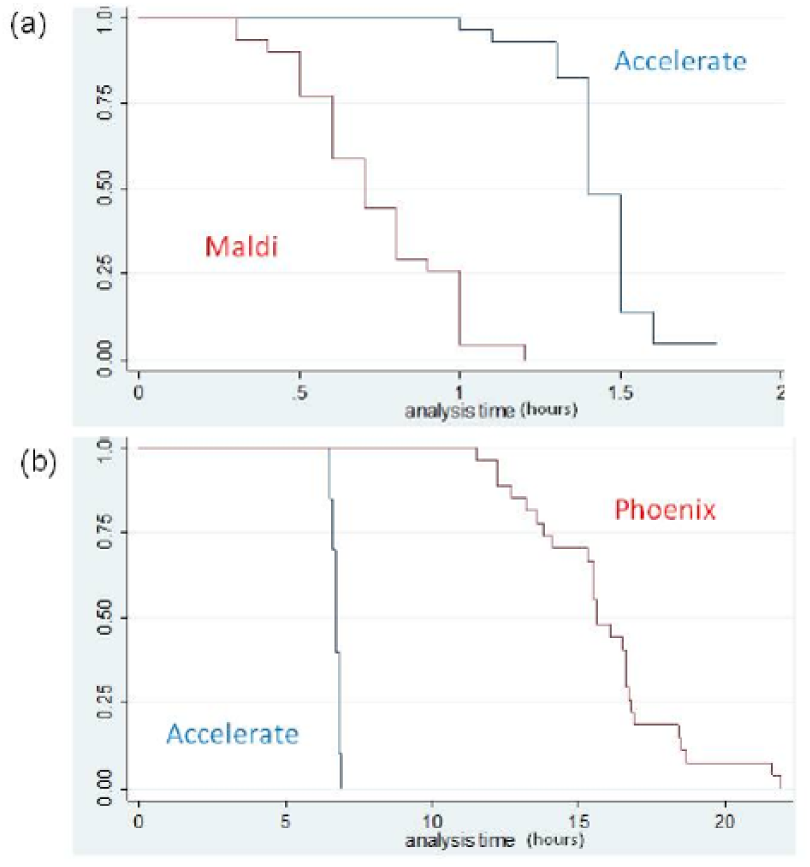
Comparison of time to identification and time to antimicrobial susceptibility testing (AST) using Accelerate Pheno™ and MALDI-TOF MS and the AST BD Phoenix system. Time to identification was compared with MALDI-TOF (Bruker, Germany) (a), and time to AST reporting was compared to Phoenix (BD, US) (b). Both MALDI-TOF and Phoenix were directly performed from the bacterial pellet obtained of positive-blood cultures after Gram stain verification. a Accelerate Pheno™

The AST results showed an overall essential agreement of 92% for the MICs and an overall category agreement of 96%. Among Gram positive isolates essential and category agreements of 100% were observed. However, for *S. agalactiae* and *S. pneumoniae* the Accelerate Pheno™ system only produced identification results but no AST. In Gram negative isolates ten discrepancies were detected, 7 major errors and 3 minor errors. In all 10 cases the reference E-test results agreed with the BD Phoenix system. The discrepancy results for AST were observed particularly in *P. aeruginosa* isolates, where the MICs estimated by the Accelerate Pheno™ system were higher than those reported by the BD Phoenix system. More specifically, the MICs results of piperacillin-tazobactam were 32 µg/ml with the Accelerate Pheno™ system and <=4 µg/ml with the BD Phoenix system for two isolates. The Accelerate Pheno™ system showed a MIC of 16 µg /ml for Ceftazidime in two strain, but with the BD Phoenix system the MIC was 2 µg/ml in both cases; the Accelerate Pheno™ system showed a MIC of 8 µg/ml and 16 µg/ml for imipenem in two strains, whereas the BD Phoenix system showed a MIC of 2 µg/mL in both cases. Finally, the Accelerate Pheno™ system presented a MIC of 8 µg/mL for gentamicin but with the BD Phoenix system the MIC result was 2 µg/mL. All discrepancies were resolved using E-Test strips which were always in agreement with the BD Phoenix system.

## Discussion

The results of this study support the idea that the Accelerate Pheno™ system may improve turn-around time for identification and AST testing for bacterial blood-stream infections. As previously reported (5–7), in some cases the Accelerate Pheno™ system detected more than one microorganism in mono-microbial infections. In addition, the ID panel failed to show any results for yeasts, which were detected in two cases by subculture. However, we used a previous version of the software; the new version of this software includes an algorithm modification to optimize its performance. A recent study established an overall sensitivity and specificity for yeasts of 83.3% and 98.4%, respectively (<u>8)</u>. The overall essential agreement and the overall category agreement in our study concurs with the multicentre study of Pancholi and colleagues which reported values of 97% (9). The AST for piperacillin-tazobactam and *P. aeruginosa* has also been optimized in the latest version of the software (5). The delay in the correct antibiotic therapy in patients with sepsis is a risk factor for mortality (10, 11). Moreover, a recent study reported shorter times from empirical to final antibiotic treatment using the Accelerate Pheno™ system when compared to conventional phenotypic methods (12). In our study none of the isolates analysed showed a specific mechanism of antibiotic resistance that required changing empirical treatment, but the Accelerate Pheno™ system is capable of detecting different resistance profiles (13,14).

In conclusion, the Accelerate Pheno™ system could improve turn-around time in the management of patients with sepsis. The reduced number of samples was a limitation in this study. Further prospective validation studies will determine its cost-efficacy and clinical effectiveness.

## AUTHOR CONTRIBUTIONS

These contributions follow the Contributor Roles Conceptualization: MFP, MA, CCP; Data curation: MFP, JM, CCP; Formal analysis: MFP, ER, AF, IAC, CCP; Funding acquisition: MA, FM, JV; Investigation: MFP, CCP; Methodology: MFP,JP, MA, FM, JV,CCP; Supervision: MA, FM, JV, CCP; Writing – review & editing: all authors.

